# Creation of X-linked Alport Syndrome Rat Model with *Col4a5* Deficiency

**DOI:** 10.1101/2021.04.13.439726

**Authors:** Masumi Namba, Tomoe Kobayashi, Mayumi Kohno, Takayuki Koyano, Takuo Hirose, Masaki Fukushima, Makoto Matsuyama

**Affiliations:** Division of Molecular Genetics, Shigei Medical Research Institute, Okayama, Japan; Division of Nephrology and Endocrinology, Faculty of Medicine, Tohoku Medical and Pharmaceutical University, Sendai, Japan; Department of Endocrinology and Applied Medicine, Tohoku University Graduate School of Medicine, Sendai, Japan; Shigei Medical Research Hospital, Okayama, Japan

## Abstract

Alport syndrome is an inherited chronic human kidney disease, characterized by glomerular basement membrane abnormalities. This disease is caused by mutations in *COL4A3, COL4A4*, or *COL4A5* gene. The knockout mice for *Col4α3, Col4α4*, and *Col4α5* are developed and well characterized for the study of Alport syndrome. However, disease progression and effects of pharmacological therapy depend on the genetic variability. This model is reliable only to mice. Therefore in this study, we created a novel Alport syndrome rat model utilizing rGONAD technology. *Col4α5* deficient rats showed hematuria, proteinuria, high levels of BUN, Cre, and then died at 18 to 28 weeks of age (Hemizygous mutant males). Histological and ultrastructural analyses displayed the abnormalities including parietal cell hyperplasia, mesangial sclerosis, and interstitial fibrosis. Then, we demonstrated that α3/α4/α5 (IV) and α5/α5/α6 (IV) chains of type IV collagen disrupted in the *Col4α5* deficient rats. Moreover, immunofluorescence analyses revealed that some glomeruli of *Col4α5* mutant rats were found to be disrupted from postnatal day 0. Thus, *Col4α5* mutant rat is a reliable candidate for Alport syndrome model for underlying the mechanism of renal diseases and further identifying potential therapeutic targets for human renal diseases.

## Introduction

Type IV collagen networks are the structural foundation for all basement membranes, consist of 6 different chains from COL4A1 to COL4A6 in mammals.^1, 2^ The 6 chains assemble into 3 types of heterotrimers (protomers), which have distinct tissue distribution and function. In the mammalian kidney, the glomerular basement membrane (GBM), requiring long-term maintenance of the glomerular filtration functions, is composed of α3/α4/α5 (IV) isoforms. Whereas α1/α1/α2 (IV) isoforms are found in all basement membranes, and α5/α5/α6 (IV) isoforms are detected in Bowman’s capsule (BC).

Alport syndrome is a basement membrane disorder, characterized by hereditary nephropathy that results in irreversible, progressive renal failure.^3^ Alport syndrome is caused by mutations in *COL4A3, COL4A4*, or *COL4A5* gene encoding type IV collagen α3, α4, or α5 chains. Defects in the *COL4A5* gene cause X-linked Alport syndrome, which accounts for about 80% of Alport syndrome.^5^ And the remaining cases are associated with mutations in *COL4A3* or *COL4A4* genes. The pathological events of Alport syndrome are very similar, are found that lack of *COL4A3, COL4A4*, or *COL4A5* resulted in the degradation of α3/α4/α5 (IV) heterotrimers.^1^

In the past decades, several Alport syndrome animal models have been produced in mice. The knockout mice for *Col4α3, Col4α4*, and *Col4α5* are developed and well characterized.^6-9^ Moreover, there are several Knock-out and Knock-in mice harboring missense, nonsense, deletion mutant of *Col4α5*.^10, 11^ These model mice led important aspects of the renal disease. Whereas, there are some significant differences depend on genetic background in Alport syndrome model mice.^8, 12, 13^ For example, 129×1/Sv or C57BL/6 was different patterns of disease progression in *Col4α3*^*-/-*^ mice.^14^ Indeed, in the human patients, the disease progression and effects of pharmacological therapy were shown to genetic variability. However, in spite of variabilities displayed in the genetic background, only a few mammal models (only mice and dogs) have been established for Alport syndrome.

The laboratory rat (*Rattus norvegicus*) is a common experimental model for the human diseases and the drug testing.^15-17^ For example, Wistar Kyoto (WKY) rat strain is known to be uniquely susceptible to crescentic glomerulonephritis among the strains tested.^18-21^ Although there are several advantages comparing with mice, production of genetically engineered rat models has not yet been extensively proceeded during the past decades. The recent CRISPR/Cas9 system is the simplest for generating rats carrying a modified genome,^22, 23^ however, the standard method for genome-editing in mammals involves 3 major steps: isolation of zygotes from females, micromanipulation *ex vivo*, and transfer into pseudopregnant females. These 3 steps require extremely high level of technical expertise and its proficiency of the researchers as well as technicians. To simplify these complexed and laborious processes, we developed more simple and reliable method for genome editing in mice and rats are now available, namely *i*-GONAD and rGONAD (rat Genome-editing via Oviductal Nucleic Acids Delivery).^24-27^ The rGONAD method involves 2 key steps: an injection of the solution containing Cas9 protein, guide RNA and single strand DNA (ssDNA) into the oviduct, followed by electroporation. The rats that have undergone rGONAD are bred in routine way until birth. Moreover, rGONAD is highly efficient in both knock-out and knock-in rats.^26^

In the present study, we attempted to create Alport syndrome rat model using rGONAD technology. We developed *Col4α5* deficient mutant rats, identical to the *Col4α5* G5X mutant mice,^9^ which showed progressive glomerular disease and CKD phenotypes with renal fibrosis. And we investigated whether the mutant rats were applicable for the Alport syndrome model.

## Results

### Generation of *Col4α5* deficient rats by the rGONAD method

On the basis of human mutations as described previously, we introduced novel *Col4α5* deficient rats with CRISPR/Cas9 and rGONAD technology.^26^ Tandem STOP codons were integrated into 27 bases after the first ATG in rat *Col4α5* gene (Supplemental Figure 1A). The mutation was verified by PCR followed by DNA sequencing, resulting to be expressed only 9 amino acids of COL4A5 at its N-terminus (Supplemental Figure 1B, C). We detected no COL4A5 protein expression in the *Col4α5* deficient male rats by immunofluorescence and Western blot analyses with monoclonal antibodies against the NC1 (C-terminus) domains of type IV collagens (See below; Figure 5, 6). In addition, the mutant rats which were integrated the other flame-shift tandem STOP codons (Col4a5 15aa stop; Supplemental Figure 2) and were deleted 56bp including the first ATG (Col4a5 56bp deletion; Supplemental Figure 3), also generated, and revealed the same phenotype as Alport syndrome rats (Supplemental Figure 2, 3). These data collectively indicate the successful generation of *Col4α5* deficient rats.

### Physiological analyses in *Col4α5* mutants

*Col4α5* deficient rats were viable, fertile, and expected Mendelian ratios. However, all hemizygous mutant males died from 18 to 28 weeks of age (n=58; Figure 1A). Heterozygous mutant females died at 35 to 100 weeks of age, among them, approximately 30 % of females survived over 100 weeks (n=33, Figure 1A). To assess of functional and histological abnormalities of the kidney, we measured the hematuria and proteinuria in *Col4α5* deficient rats. Hematuria was found from postnatal 21 days in hemizygous mutant males, and from 4 weeks of age in heterozygous mutant females (data not shown). Proteinuria was observed (about 6.0 mg/16 hours) by 6 weeks of age in hemizygous mutant males (see Supplemental Table 1), then, increased > 20.0 mg/16 hours in all of mutant males after 12 weeks of age (n=35), and 62% of mutant females in 16 week of age (n=28/45) (Figure 1B). On the contrary, these phenomena were not detected in wildtype male and female rats in any week of age (Figure 1B).

**Figure 1.**
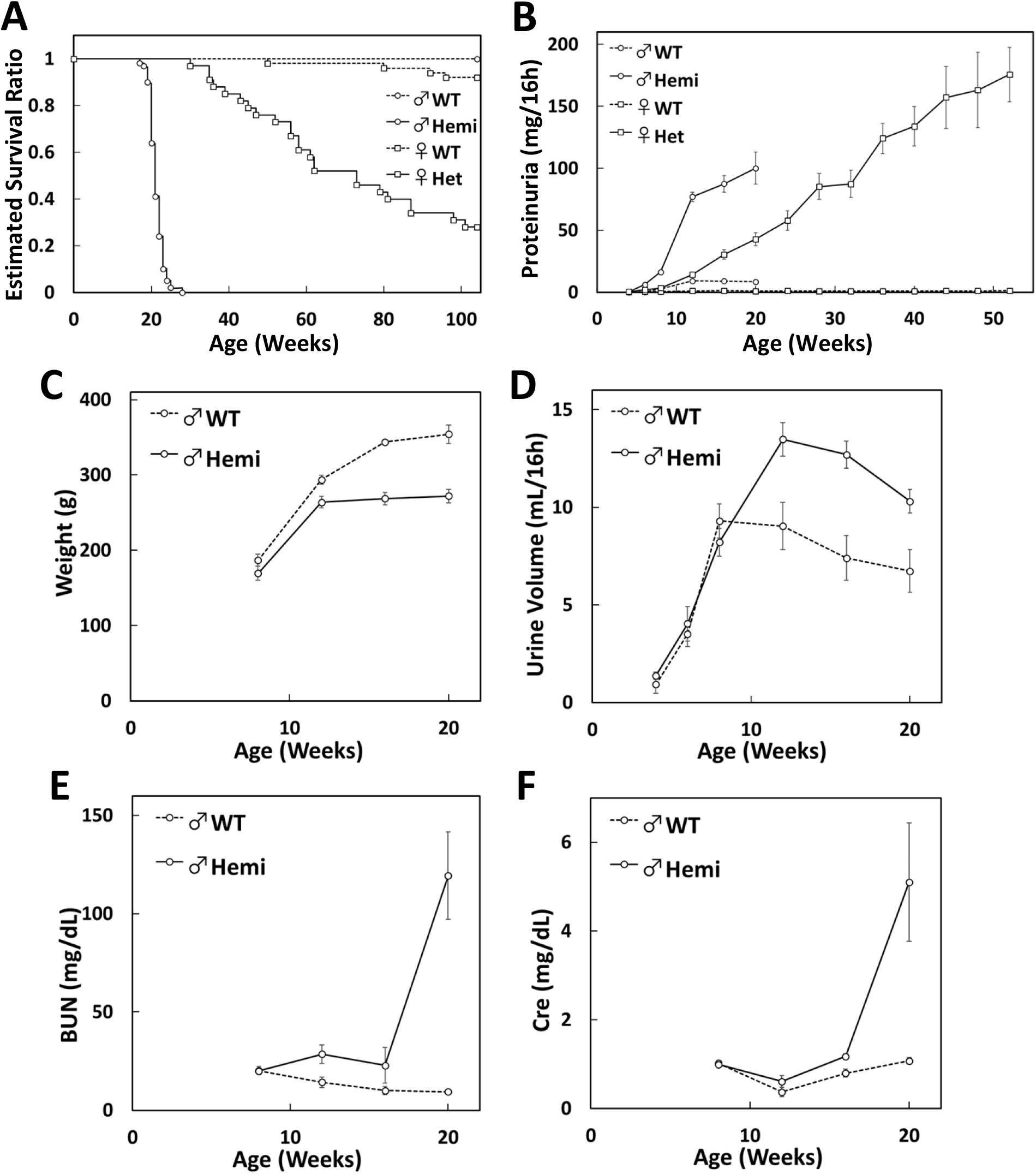
Physiological analyses in *Col4α5* mutant rats. (A) Estimated survival functions in wildtype (WT) and *Col4α5* mutant (Hemi; hemizygous males, het; heterozygous females) rats. (B) Proteinuria in wildtype and *Col4α5* deficient rats from 4 to 52 weeks of age. (C-F) Measurements of body weight (C), urine volume (D), blood urea nitrogen (BUN) of serum (E), and serum Creatinine (Cre) in wildtype and *Col4α5* mutant males from 4 to 20 weeks of age.

Mutant males showed decrease of body weight and increase of urine volume, compared with those of wildtype males (Figure 1C, D). The levels of blood urea nitrogen (BUN) and serum creatinine (Cre) in serum highly increased in 20 weeks of age of mutant males (Figure 1E, F).

### Renal histology in *Col4α5* deficient rats

The kidney sections from wildtype and *Col4α5* deficient rats were stained with hematoxylin and eosin (HE), Periodic acid Schiff (PAS), periodic acid methenamine silver (PAM), and Masson trichrome (MT) (Figure 2). At 8 weeks of age, the glomeruli exhibited capillary tuft collapse in hemizygous mutant males (Figure 2A). However, the kidney displayed overall sparing of the tubulointerstitium (Figure 2A). By the 20 weeks of age in hemizygous mutant males, substantial numbers of glomeruli revealed the abnormalities, including parietal cell hyperplasia mimicking crescent formation, focal sclerosis, and with overall sparing of the tubulointerstitium (Figure 2B, Supplemental Figure 4). The kidney sections from heterozygous mutant females at 8-20 weeks of age displayed focal abnormalities of the glomeruli and tubulointerstitium (Figure 2A, B, Supplemental Figure 4).

**Figure 2.**
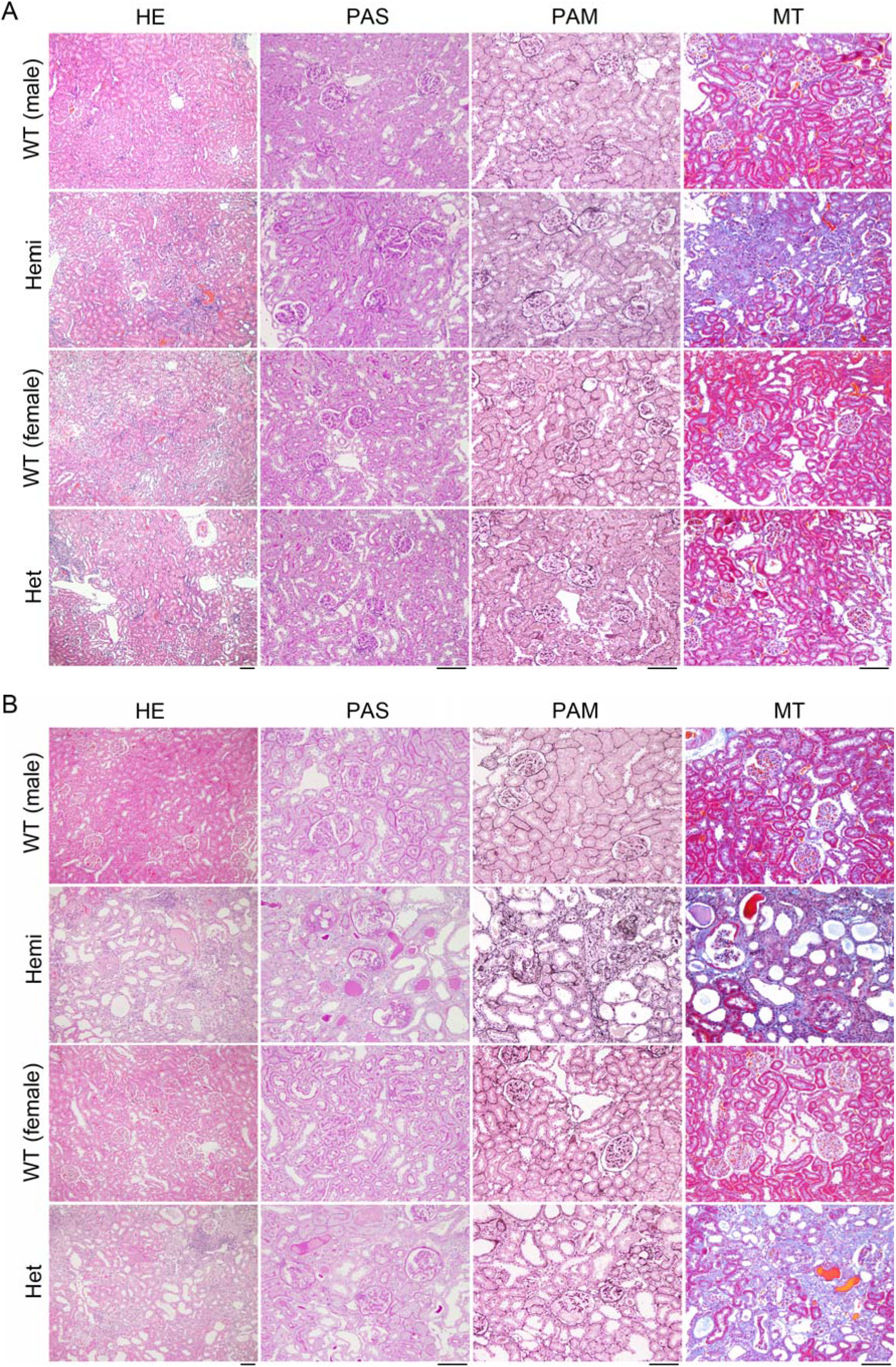
Histological analyses of the *Col4α5* deficient kidneys. Representative microscopic images in wildtype (WT) and *Col4α5* mutant (Hemi; hemizygous males, Het; heterozygous females) rats at 8weeks (A) and 20 weeks (B) of age. These tissue sections were prepared and stained with hematoxylin and eosin (HE), Periodic acid Schiff (PAS), periodic acid methenamine silver (PAM), and Masson trichrome (MT). *Scale bars*, 100 µm.

The 3 dimensional ultrastructure of GBM observed by low-vacuum scanning electron microscopy (LVSEM) revealed the coarse meshwork structure of GBM with numerous pin-holes in hemizygous mutant males, in contrast to the smoothly arranged surface in the wildtype males (Figure 3, Supplemental Figure 5).

**Figure 3.**
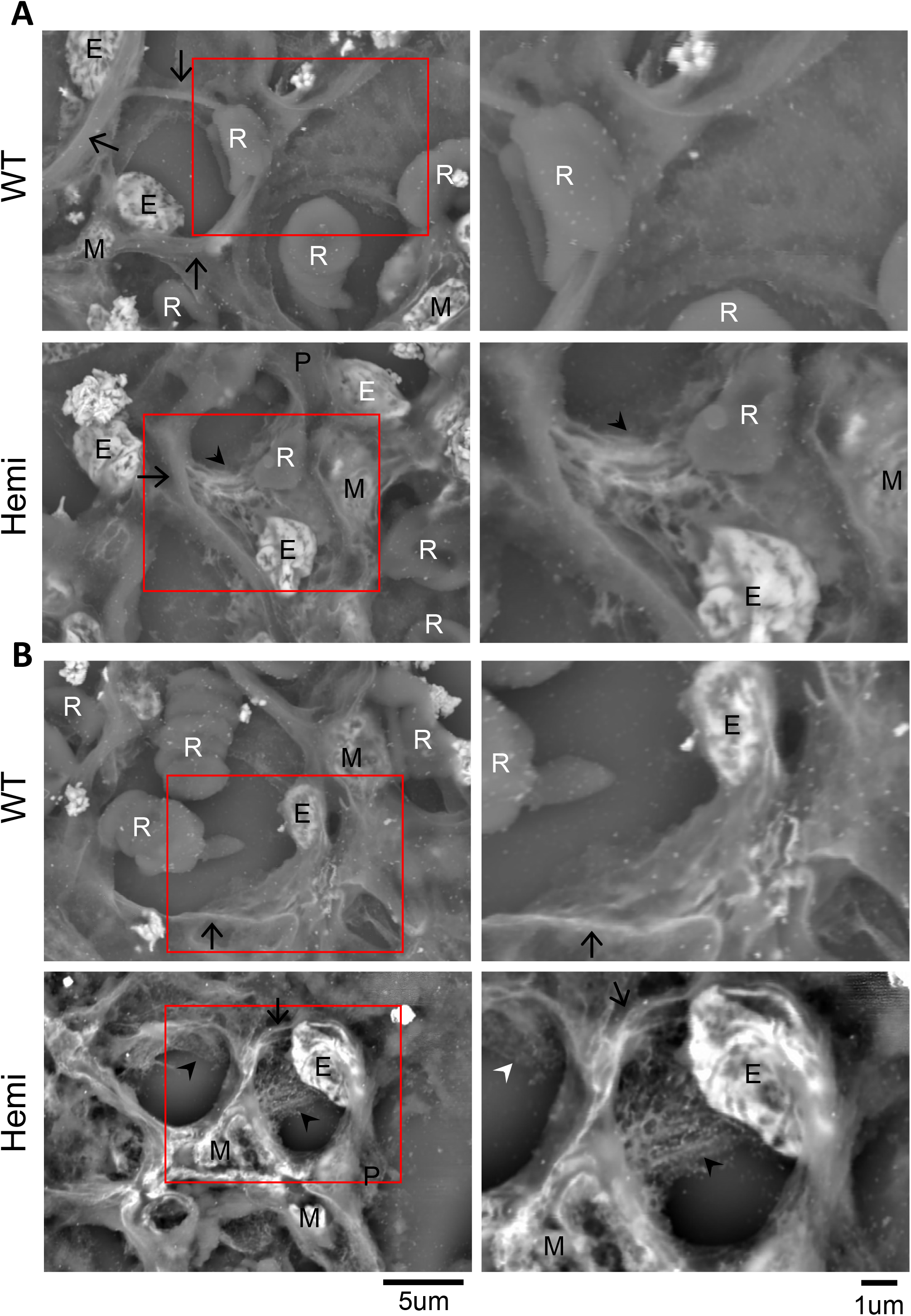
Electron photomicrographs of glomerular basement membranes in *Col4α5* mutant rats. Representative Low-vacuum scanning electron microscopy (LVSEM) images in wildtype (WT) and *Col4α5* mutant (Hemi) males at 8weeks (A) and 20 weeks (B) of age. Arrowheads indicate the coarse meshwork structure of the GBMs. Arrows also indicate cut side of the capillary walls. Red insets are revealed the higher magnification of left panels. E: Endothelial cells, M: Mesangial cells, P: Podocytes, R: Red blood cells. *Scale bars*, 5 µm (left), 1 µm (right).

### Renal glomerular and tubulointerstitial fibrosis in *Col4α5* deficient rats

To evaluate the fibrosis of glomeruli and tubulointerstitium in *Col4α5* deficient rats, we examined the kidney sections from wildtype and *Col4α5* deficient rats stained with antibodies specific for α-smooth muscle actin (α-SMA) and fibronectin (Figure 4). Immunostaining showed expression level of α-SMA, a maker of renal glomerular and tubulointerstitial fibrosis, was increased around the glomeruli in the kidneys from hemizygous mutant males when compared with those of wildtype littermates at 20 weeks of age, but was not detected in the renal tubular epithelia (Figure 4A). The kidney sections from the heterozygous females were rarely detected by 20 weeks of age (Figure 4A, Supplemental Figure 6). In contrast, the expression level of fibronectin, a marker of pathological deposition of extracellular matrix (ECM), was increased in 12 weeks of age of the hemizygous mutant males (Figure 4B). We then observed fibronectin expression in the heterozygous mutant females at 12 weeks of age (Figure 4B, Supplemental Figure 7). These results suggest that the increase of fibronectin expression might represent the mesangial sclerosis and interstitial fibrosis in *Col4α5* deficient males and females.

**Figure 4.**
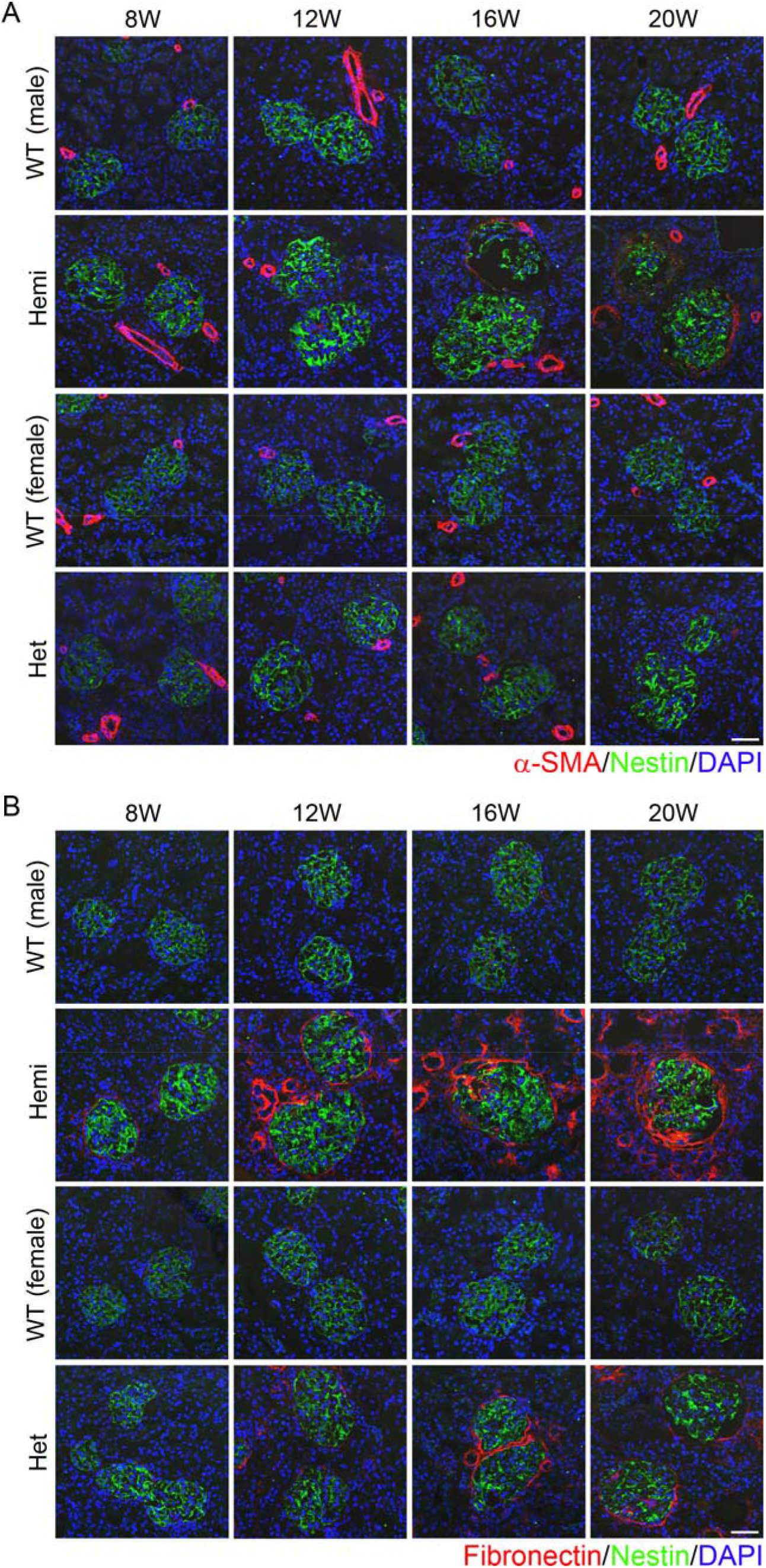
Renal fibrosis in *Col4α5* deficient rats. (A, B) Immunostaining of kidney sections with αSMA (A) or fibronectin (B) (red), nestin (green; glomeruli), and DAPI (blue; nuclei) in wildtype (WT) and *Col4α5* mutant (Hemi; hemizygous males, Het; heterozygous females) rats from 8 to 20 weeks of age. *Scale bars*, 50 µm.

### The expression of type IV Collagen α1-6 in *Col4α5* deficient rats

The type IV collagen networks comprised of α1/α1/α2 (IV) (all basement membranes), α3/α4/α5 (IV) (GBM), and α5/α5/α6 (IV) (Bowman’s capsule) protomers were observed in the glomeruli. To examine whether localizations and/or the levels of the proteins were changed in the *Col4α5* deficient rats, we investigated *Col4α5* deficient rats with graded levels of the other’s type IV collagen α1, α2, α3, α4, and α6 (IV). First, we produced novel monoclonal antibodies that specifically recognize rat COL4A6 protein. We obtained rCol4A6 antibodies by injecting the antigen in *Col4a5* deficient males, but we could not obtain the specific antibody injected to wildtype rats at all. Immunoblot analysis revealed that the monoclonal anti-rCOL4A6, reacted specifically with a band corresponding to the position of the similar molecular weights in recombinant rCOL4A6 proteins, and only reacted with rCOL4A6, but not with other rCOL4 proteins (Supplemental Figure 8).

To determine whether localization of type IV collagens of α1-6 (IV) in the kidney of *Col4α5* deficient rats changed, we performed the immunofluorescence analyses. At 8-20 weeks of age, COL4A5 expression was absent in the hemizygous mutant males, and present in a mosaic pattern in heterozygous mutant females (Figure 5A, B, Supplemental Figure 9-12). The expressions of COL4A3, COL4A4, and COL4A6 were also absent in *Col4α5* deficient males, and present in a mosaic pattern in heterozygous mutant females (Figure 5A, B, Supplemental Figure 9-12). At 8 weeks of age, in contrast, expressions of COL4A1 and COL4A2 were present in *Col4α5* deficient rats (Figure 5A, Supplemental Figure 9). At 20 weeks of age, the kidney showed a strong accumulation of COL4A1 and COL4A2 proteins in both GBM and Bowman’s capsule (Figure 5B, Supplemental Figure 9-12). These data suggest that α3/α4/α5 (IV) and α5/α5/α6 (IV) chains of type IV collagen disrupted in the *Col4α5* deficient rats.

**Figure 5.**
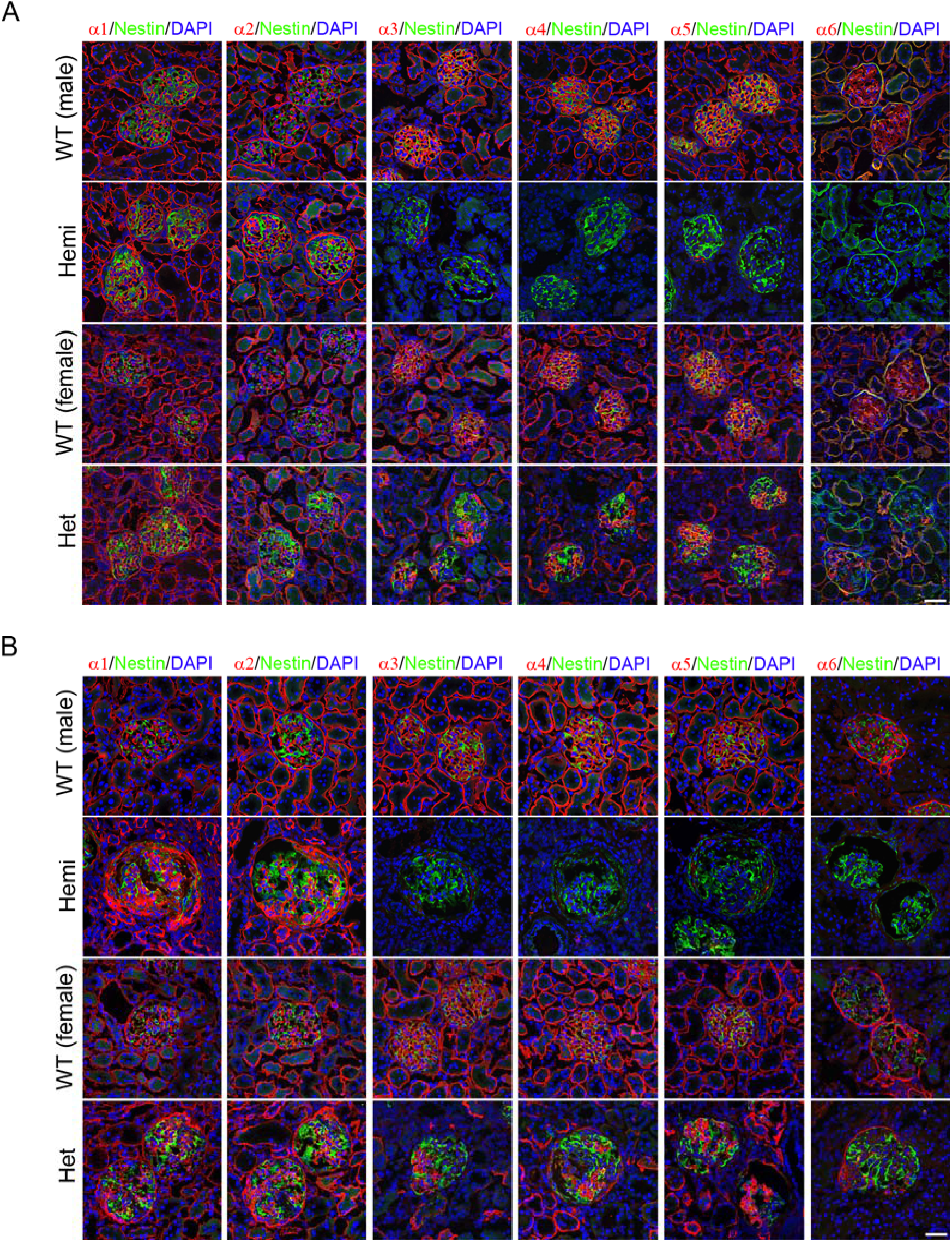
Type IV collagen distributions in the *Col4α5* deficient kidneys. (A, B) Immunofluorescence analyses of kidney sections with antibodies against α1-6 (IV) (red), nestin (green; glomeruli), and DAPI (blue; nuclei) in wildtype (WT) and *Col4α5* mutant (Hemi; hemizygous males, Het; heterozygous females) rats at 8weeks (A) and 20 weeks (B) of age. *Scale bars*, 50 µm.

To verify the results of the immunostaining, we analyzed protein expressions noncollagenous domains 1 (NC1) that were prepared from kidneys of wildtype and hemizygous mutant males by Western blot analyses (Figure 6). There were no bands detectable NC1 domains of type IV Collagen α3, α4, and α5 (IV), derived from α3/α4/α5 (IV) complexes, in kidney of *Col4α5* deficient males (Figure 6). Then, the NC1 domains of type IV Collagen α5 and α6 (IV), derived from α5/α5/α6 (IV) complexes, were also absent, whereas the blotting for the NC1 domains of type IV Collagen α1 and α2 (IV), derived from α1/α1/α2 (IV) complexes, were present in the hemizygous mutant males as well as wild type (Figure 6). Therefore, these findings fully corroborate the results of immunofluorescence analyses.

**Figure 6.**
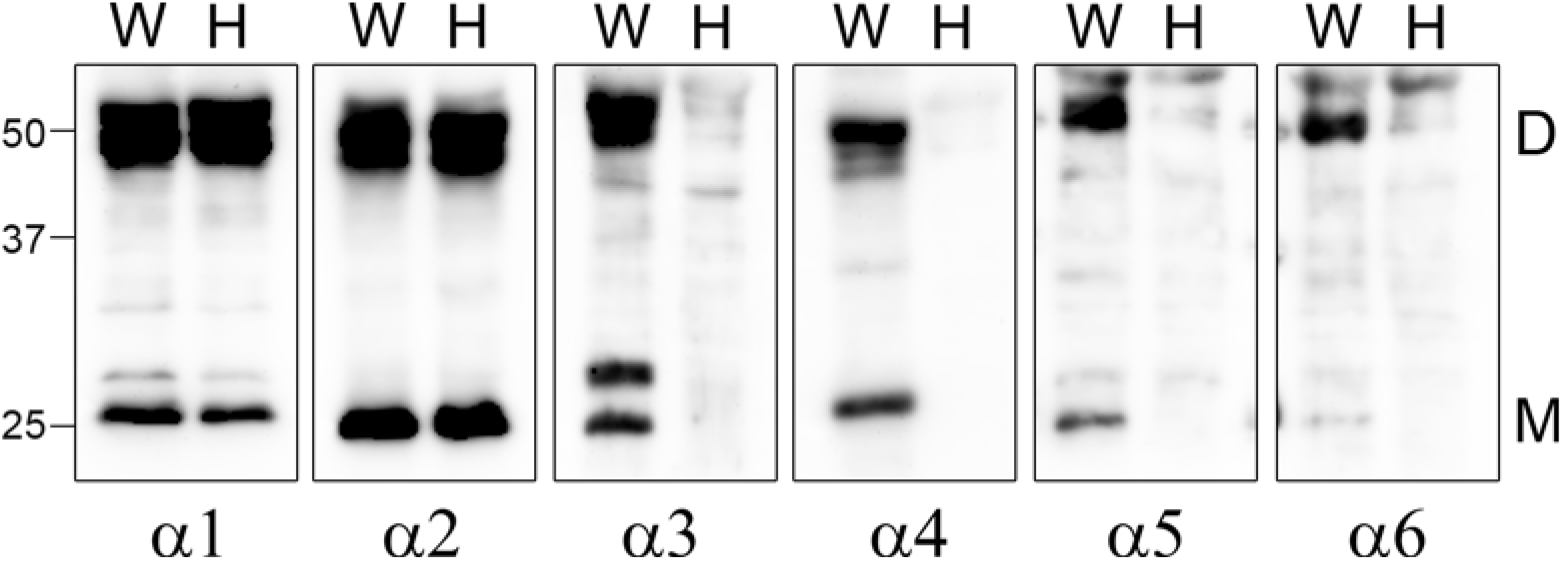
Western blot analyses of type IV collagen in the *Col4α5* mutant kidneys. Collagenase-solubilized renal basement membranes from wildtype (W) and *Col4α5* mutant (H) males at 8 weeks of age were separated by SDS-PAGE and blotted with antibodies that the specifically recognized α1-6 (IV) NC1 domains. D: NC1 dimer, M: NC1 monomer.

### *Col4α5* deficient rats in early renal developmental stage

Alport syndrome has well been studied in terms of adult renal pathogenesis, although little is still known in the early development. We focused to investigate early renal pathology with the *Col4α5* deficient rats. To determine when COL4A5 protein expresses in the postnatal kidney, we performed immunostaining on wildtype and on *Col4α5* deficient males from birth to 28 days of age (Figure 7A). After birth (P0), COL4A5 was expressed in only GBM in wildtype rats (Figure 7A). During postnatal day (P7), this expression was spread to Bowman’s capsule and basement membrane of ureter epithelial cells (Figure 7A). There were no COL4a5 staining detectable in the *Col4α5* mutant male kidneys with anti-COL4A5 antibody (Figure 7A).

**Figure 7.**
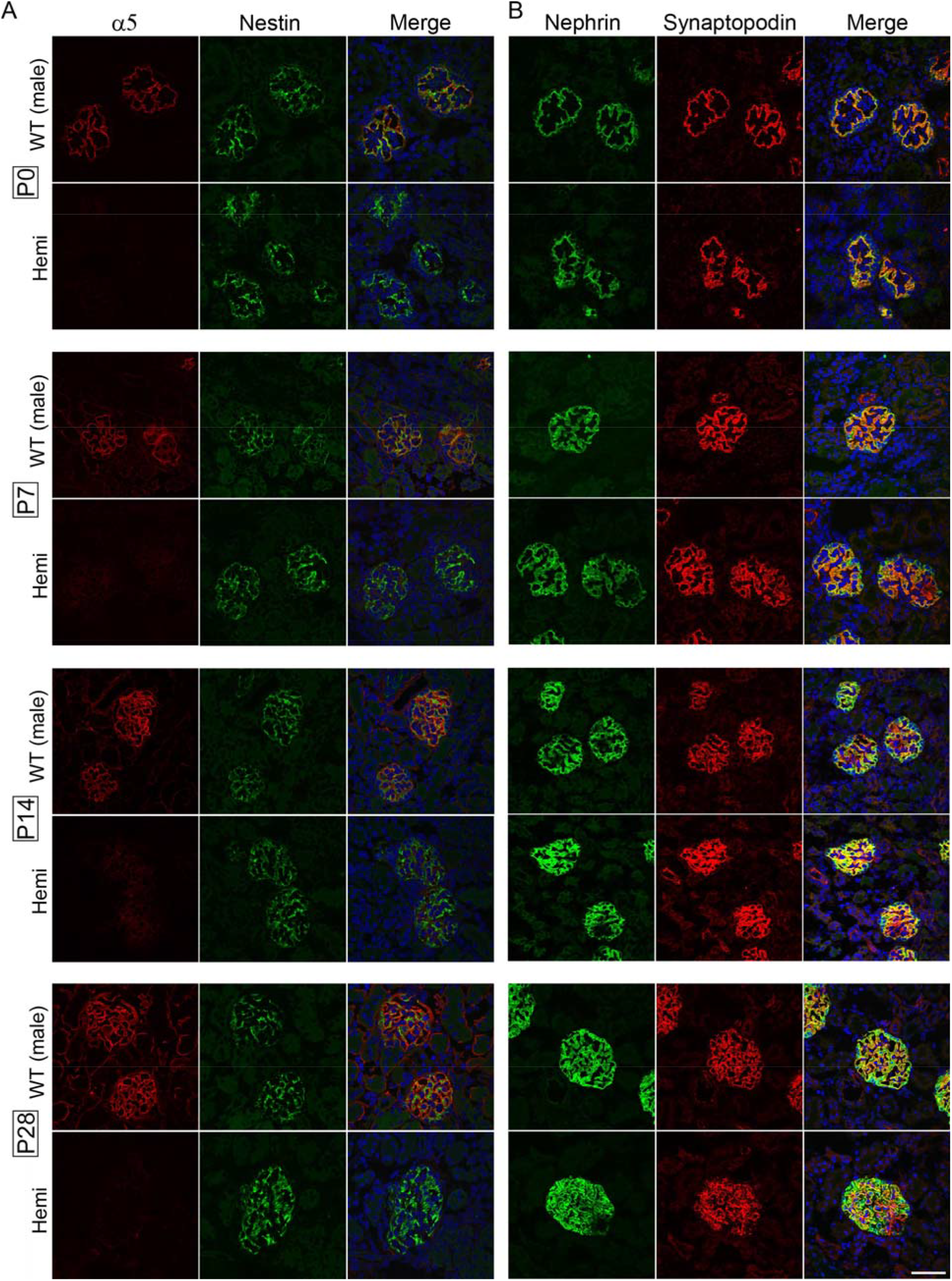
Early renal pathology with *Col4α5* deficient rats. (A, B) Immunofluorescence analyses of the rat kidney sections with antibodies against (A): α5 (IV) (red), nestin (green; glomeruli), and DAPI (blue; nuclei), (B): Synaptopodin (red), nephrin (green), and DAPI (blue; nuclei), in wildtype (WT) and *Col4α5* mutant (Hemi) male rats after birth, postnatal (P) 7, 14, and 28 days of age. *Scale bars*, 50 µm.

To detect renal failure in postnatal stage, we examined expression of nestin, nephrin, and synaptopodin in *Col4α5* hemizygous males. Immunofluorescence analyses showed nestin, a marker of intermediate filament protein, was located orderly in glomeruli of wildtype kidneys after birth (Figure 7A). In contrast, the expression of some glomeruli was found to be disrupted from postnatal day 0 (Figure 7A). Immunofluorescence analyses also revealed that expression of nephrin, a marker of transmembrane cell adhesion molecule, was found to locate at slit diaphragm in GBM, and synaptopodin, a marker of slit diaphragm proteins expressed in GBM, were disorganized, and found aggregated (Figure 7B). Thus, these observations certainly suggest that slit diaphragm in GBM was already disrupted at the postnatal development of the *Col4α5* deficient kidneys.

## Discussion

The present study showed that we successfully created a novel Alport syndrome rat model with rGONAD technology. *Col4α5* deficient rats revealed typical physiological, pathological, and also histological characteristics of Alport syndrome. Furthermore, it will be especially useful for studying progress of occurrence of the renal diseases of Alport syndrome beginning from postnatal stages.

Occurrence of Alport syndrome is caused by mutations in *COL4A3, COL4A4*, or *COL4A5*.^4^ Several mouse models in *Col4α3, Col4α4*, and *Col4α5* mutation have been known and characterized.^6-10, 13^ *Col4α3*^*-/-*^ or *Col4α4*^*-/-*^ mice are well used for studying treatments of Alport syndrome compared to *Col4α5*^*-/Y*^ mice.^28-31^ Because, a survival ratio of *Col4α5*^*-/Y*^ mice was so diffusive ranged from 6 to 34 weeks,^9^ in contrast, those of *Col4α3*^*-/-*^ or *Col4α4*^*-/-*^ mice were more uniform (from 13 to 26 weeks).^32, 33^ In this study, *Col4α5* mutant males died at 18 to 28 weeks (about 95 % from 20 to 26 weeks) of age. Thus, it is useful to study experiments for prolong the lifespan of *Col4α5* deficient rats for effects of the drug, food, and other factors.

In the present study, we were able to produce *Col4α5* mutant rats of WKY strain. Some number of glomeruli exhibited capillary tuft collapse at 8 weeks of age, and more glomeruli revealed abnormalities with crescent formation and focal sclerosis at the 20 weeks of age in *Col4α5* mutant males. The levels of BUN and Cre in serum highly increased in 20 weeks of age, and then died at around 24 weeks of age in hemizygous mutant males. These physiological and pathological features correspond to those of previously reported *Col4α3*^*-/-*^ with C57BL/6 mouse strain.^12, 14^ However, *Col4α3*^*-/-*^ with 129×1/Sv mouse strains showed much earlier progression of disease, died at around 12 weeks of age.^12, 32, 34^ Therefore, comparison of a survival ratio between different strains of mice and rats should be carefully studied them, and further studies are required to draw a definite conclusion. In further study, other rat strains of *Col4α5* mutant using rGONAD method can be expected.

The laboratory rat has long been recognized as a preferred experimental animal in wide areas of biomedical science.^22, 35^ In some situation, rat is considered as a more relevant model among mammals. For example, physiology of rat is greatly well documented, because its larger body size affords the opportunity for serial blood draw studies. In particular, blood pressure measurement by telemetry is easier to perform and more reliable in rats compared to smaller mice.^36, 37^ Moreover, there are several rat strains in order to perform specific studies, i,e, SHR for hypertension.^38^ Although genome engineering experiments of rat have yet not been extensively proceeded, it is readily applicable to produce the gene-editing mutant rats in *Col4α5* gene of the strains by rGONAD method. The model rats might provide new insight into the study of the relationship between renal disease such as Alport syndrome and other blood pressure related diseases.

In conclusion, we have described creation of a novel rat model of Alport syndrome. This rat model should be available for the study of progressive renal failure having basement membrane abnormality after birth. Thus, *Col4α5* mutant rat is a reliable candidate for Alport syndrome model animal for underlying the mechanism of renal diseases and further identifying potential therapeutic targets for human renal diseases.

## Materials and Methods

### Animals

WKY/NCrl rats were obtained from Charles River. The rats were kept with a 12:12-h light: dark cycle. They were given free access to drinking water and food. All animals were handled in strict accordance with good animal practice as defined by the relevant national and/or local animal welfare bodies, and all animal works were approved by the appropriate committee (permission number: #17006).

### Generation of type IV Collagen α5 KO rats

Type IV Collagen α5 deficient rats were produced by the rGONAD method as previously described.^26^ Briefly, gene targeting strategy is designed to integrate tandem STOP codons into 27 bases after the first ATG in rat *Col4α5* gene (Supplemental figure S1A). Guide RNAs were designed using CHOPCHOP (https://chopchop.cbu.uib.no/) (Supplemental figure S1A).

For the preparation of CRISPR/Cas9 reagents, Alt-R™ CRISPR-Cas9 system (Integrated DNA Technologies [IDT, Coralville, IA]) was used in accordance with the manufacturer’s protocol. Approximately 2-2.5µl of electroporation solution was injected into the rat oviductal lumen from up-stream of ampulla using a micropipette. The electroporation was performed using a NEPA21 (NEPA GENE Co. Ltd., Chiba, Japan).

### Genotyping

Gene alteration was certified by PCR followed by DNA sequencing, as described previously.^39^ Rat genomic DNA was isolated from ear-piece or tail. Genotyping was performed by PCR with the following primers (see also Supplemental figure S1A): rCol4a5-fw, (5’-GCTCTCTTCCCAATAACCCCT-3’), rCol4a5-rv, (5’-CAATTTTGACTTCCCTGGCCA-3’).

### Urine and blood parameters

Urine samples were collected for 16 hours by placed rats in metabolic cages individually, every 4 weeks over 52 weeks (see Supplemental Table 1). Proteinuria level was measured by a modified method using 3 % sulfosalicylic acid. Blood samples (n=6 each) were collected from the tail and centrifuged at 3000 g for 5min to obtain blood serum. Blood urea nitrogen (BUN) of serum was measured using Colorimetric Detection Kit (Arbor Assays, Michigan, USA). Serum Creatinine (Cre) levels were measured Jaffe’s method (FUJIFILM Wako Pure Chemical Corporation, Osaka, Japan). All measurements were carried out according to manufacturer’s recommended protocols.

### Histological analyses

Rat kidneys were soaked in 10% buffered neutral formalin at least overnight, and embedded in paraffin after dehydrated. The embedded kidneys were sliced 1-2 µm thickness. These slides were stained with hematoxylin and eosin (HE), periodic acid Schiff (PAS), periodic acid methenamine silver (PAM) and masson’s trichrome (MT) by standard methods.

### Low vacuum scanning electron microscopy (LVSEM)

Under LVSEM, 3 dimensional ultrastructure of GBMs of Alport syndrome, the kidney was examined as described previously.^40^ In brief, renal paraffin sections of 4 µm thickness were stained with PAM. The sections on the slides were directly observed without a cover clip, with LVSEM (Hitachi TM4000; Hitachi Co. Ltd., Tokyo, Japan) at acceleration voltage of 15 kV with 30Pa.

### Preparation of Recombinant Proteins

His-tagged (for production of antibody) and MBP-tagged (for immunoblotting) NC1 domains of type IV Collagen α1-6 (V) were expressed in BL21-CodonPlus-RP (Agilent Technologies, Santa Clara, CA) transformed with pET-28a (Invitrogen) and pMAL (New England Biolabs, Beverly, MA), respectively. Each His or MBP fusion protein was purified through affinity chromatography with TALON metal affinity resin (Clonetech, Palo Alto, CA) or with amylose resin (New England Biolabs), respectively.

### Antibody

We produced rat monoclonal rCOL4A6, as described previously.^41, 42^ Briefly, the antigen emulsion was injected to *Col4a5* deficient males. The treated rats were sacrificed 21 days after the injection, and the lymphocytes were fused with SP2/0-Ag14 myeloma cells. After the cell fusion, culture supernatants were screened to confirm positive clones by solid-phase enzyme-linked immunosorbent assay (ELISA).

The following primary antibodies were used: rat monoclonal type IV collagen α1(IV) (H11), α2 (IV) (H22), α3 (IV) (H31), α4 (IV) (H43), α5(IV) (H52), (In our institute),^43^ SMA (1A4,Cell Signaling Technology, Beverly, MA); Fibronectin (ab6328, Abcam, Cambridge, UK); Nephrin (GP-N2, Progen, Germany); Nestin (66259-1-Ig); Synaptopodin (21064-1-AP, proteintech, Illinois), and MBP (New England Biolab) Primary antibodies were detected using species-specific secondary antibodies conjugated to either Alexa Fluor 488 or 555 (Molecular probes).

### Tissue extract preparation and immunoblotting

Rat kidneys were homogenized and incubated at 37°C for 4 hours with 1mg kidney lysates and 200 µg (1000 U) of collagenase (Brightase-C; Nippi, Okayama, Japan) in digestion buffer (20 mM HEPES, 10 mM CaCl_2_) with protease inhibitor cocktail (Nacalai, Kyoto, Japan). Protein concentrations for cell extracts were determined by the Coomassie Brilliant Blue staining by SDS-PAGE gels. The lysates were loaded, transferred, and subjected to Western blotting with specific antibodies as described previously.^44^

### Immunofluorescence

Immunofluorescence analyses were examined, as described previously.^45^ Briefly, rat kidneys were immersed in OCT compound, and snap-frozen in liquid nitrogen vapor. Subsequently, kidneys were sliced 5 µm cryostat sections and placed on slides. After dehydration by acetone, blocked in 5 % (v/v) donkey serum for 30 min at room temperature. Sections were then incubated with primary antibodies overnight at 4 °C followed by PBS wash and incubated with appropriate secondary antibodies for 1h at room temperature. DNA was also stained with 1 µg/ml DAPI. Fluorescence images were obtained by confocal microscopy (FV1200, Olympus, Japan).

## Abbreviations

GONAD: Oviductal Nucleic Acids Delivery
*i-*GONAD: improved GONAD
rGONAD: Rat improved GONAD
WKY: Wistar Kyoto
CRISPR: Clustered Regularly Interspaced Short Palindromic Repeats
Cas: CRISPR-associated
GBM: glomerular basement membrane
BC: Bowman’s capsule
ssDNA: single strand DNA
BUN: blood urea nitrogen
Cre: serum creatinine
HE: hematoxylin and eosin
PAS: Periodic acid Schiff
PAM: periodic acid methenamine silver
MT: Masson trichrome
LVSEM: low-vacuum scanning electron microscopy
ECM: extracellular matrix
NC1: noncollagenous domains 1,

## Ethics approval and consent to participate

All animals were handled in strict accordance with good animal practice as defined by the relevant national and/or local animal welfare bodies, and all animal works were approved by the appropriate committee.

## Authors’ contributions

MN, TK, MK, TK, MF, and MM conceived and designed the experiments, MN, TK, MK, TK, TH, and MM performed experiments, MN and MM wrote the manuscript. All authors read and approved the final manuscript.

## Acknowledgements

We are grateful to Chieko Takahashi for animal breeding, Dr. Yoshikazu Sado for technical assistance, and Dr. Tohru Okigaki for critical comments on the manuscript. We would like to thank Dr. Fumihiro Shigei, Chairman of the Board, for financial support and continuous encouragements. This work was also supported in part by the Ryobi Teien Memory Foundation and Wesco Scientific Promotion Foundation.

## Supplementary Information

Contents:

Supplemental Figure 1. Production of *Col4α5* mutant rats Supplemental Figure 2. Analyses of “Col4α 15aa stop” mutant rats

Supplemental Figure 3. Analyses of “*Col4α5* 56bp deletion” mutant rats Supplemental Figure 4. Histological analyses of the *Col4α5* deficient kidneys

Supplemental Figure 5. Electron photomicrographs of glomerular basement membranes in *Col4α5* mutant rats

Supplemental Figure 6. Renal fibrosis in *Col4α5* deficient rats Supplemental Figure 7. Renal fibrosis in *Col4α5* deficient rats

Supplemental Figure 8. Characterization of an antibody specifically recognized type IV collagen

Supplemental Figure 9. Type IV collagen distributions in the *Col4α5* deficient kidneys at 8 weeks of age

Supplemental Figure 10. Type IV collagen distributions in the *Col4α5* deficient kidneys at 12 weeks of age

Supplemental Figure 11. Type IV collagen distributions in the *Col4α5* deficient kidneys at 16 weeks of age

Supplemental Figure 12. Type IV collagen distributions in the *Col4α5* deficient kidneys at 20 weeks of age

Supplemental Table 1. Proteinuria in *Col4a5* deficient rats

