## supplemental information for "Creation of X-linked Alport Syndrome Rat Model with *Col4a5* Deficiency"

Figure S1

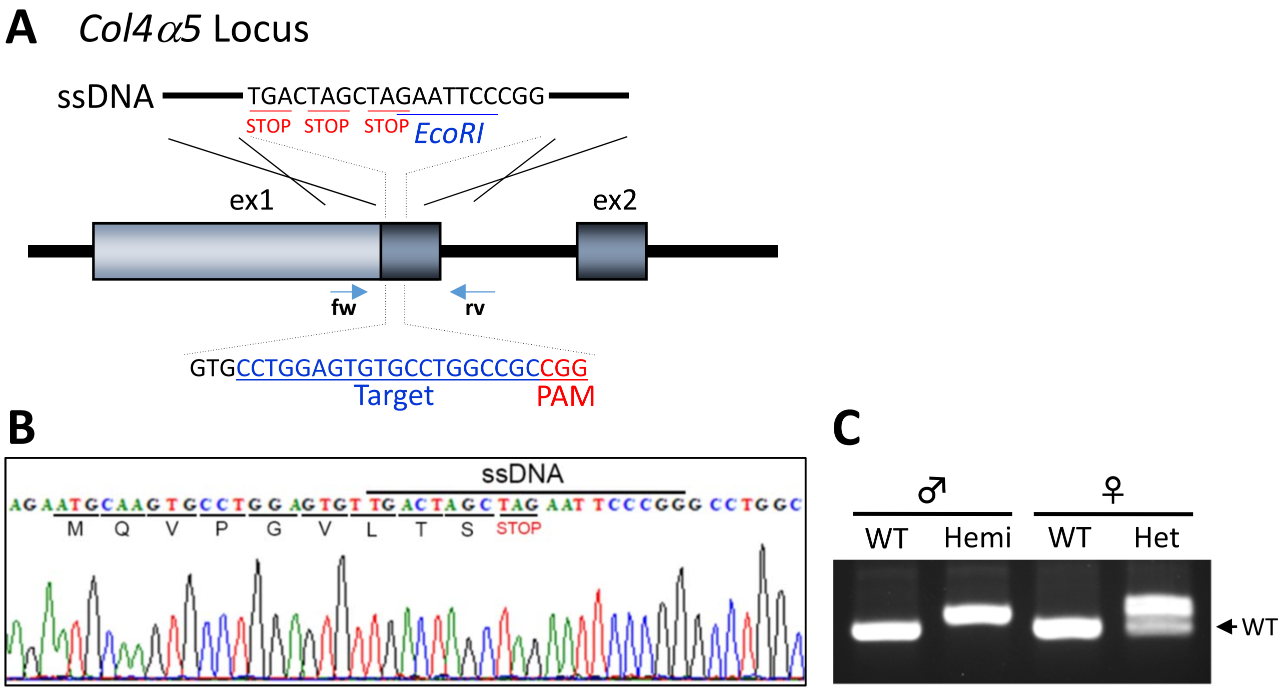


D

| Stain | Treated female | pregnant mice | Newborns | modified allele | deletion | ssODN insertion |
|---|---|---|---|---|---|---|
| WKY | 16 | 14 | 100 | 37 (37%) | 29 (29%) | 8 (8%) |

**Supplemental Figure 1. Production of *Col4α5* mutant rats**
(A) Schematic diagram of the target sequence, PAM, and ssDNA at *Col4α5* gene locus. (B) Direct sequence of mutation on *Col4α5* mutant males. (C) PCR genotyping in *Col4α5* deficient rats. (D) Efficiencies of rat *Col4α5* gene editing with rGONAD technology.

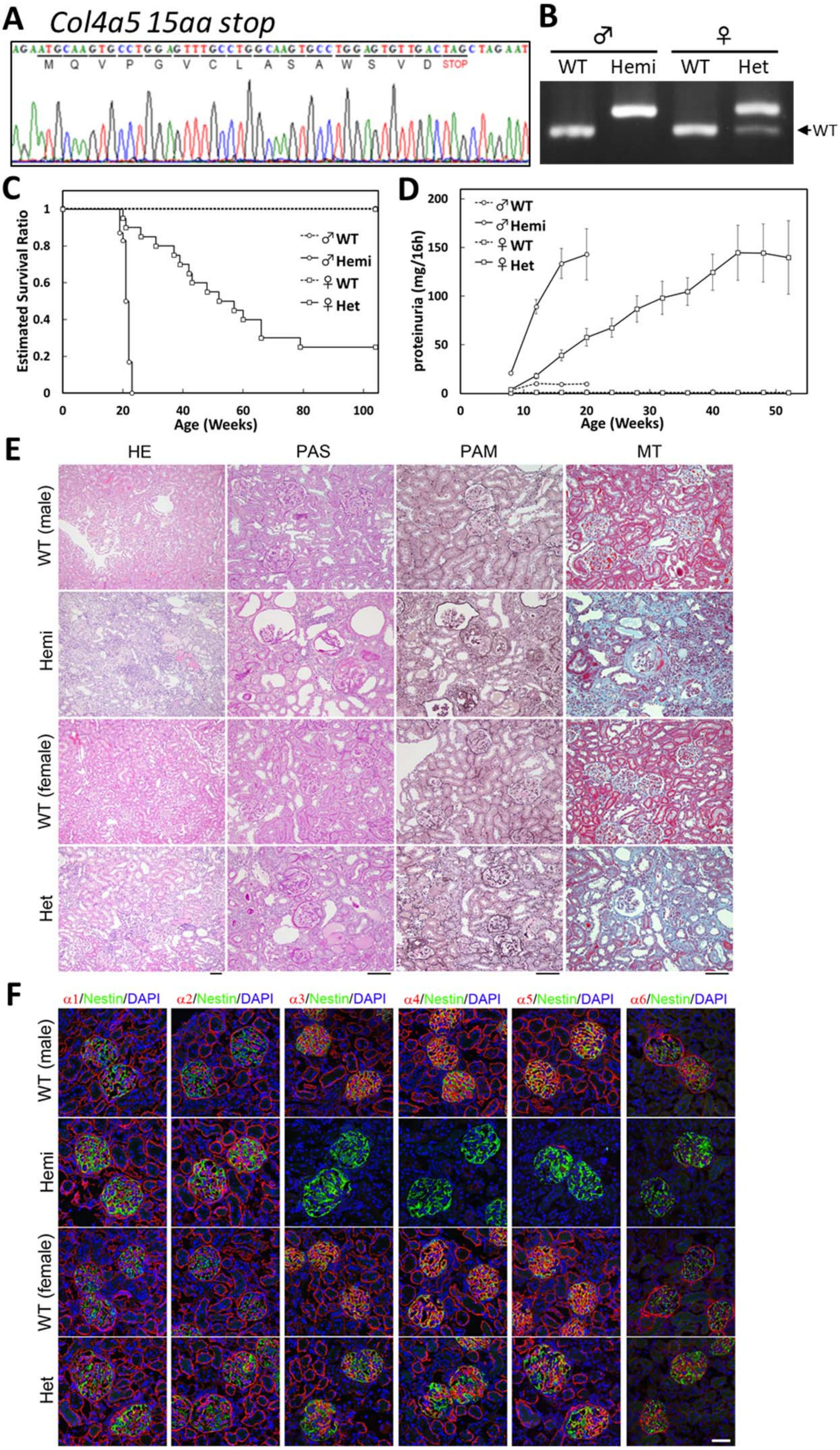


**Supplemental Figure 2. Analyses of "Col4*α* 15aa stop" mutant rats**

(A) Direct sequence of mutation. (B) PCR genotyping. (C) Estimated survival functions. (D) Proteinuria (E) Histological analyses at 20 weeks of age. (F) Type IV collagen distributions at 8 weeks of age. *Scale bars*, (E): 100 μm, (F): 50 μm.

Figure S3

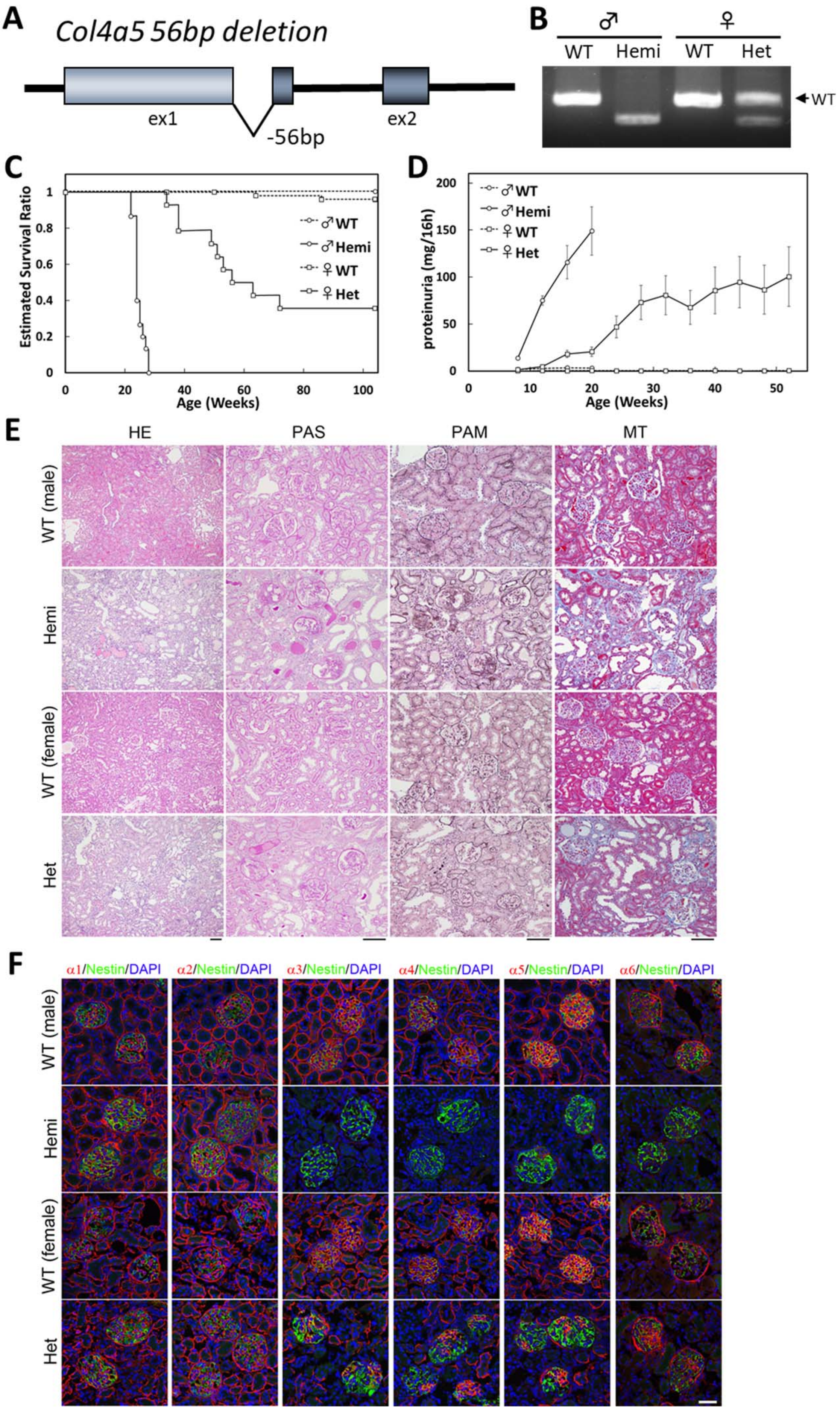


**Supplemental Figure 3. Analyses of "*Col4α5* 56bp deletion" mutant rats**
(A) Schematic diagram of the deletion mutant. (B) PCR genotyping. (C) Estimated survival functions. (D) Proteinuria (E) Histological analyses at 20 weeks of age. (F) Type IV collagen distributions at 8 weeks of age. *Scale bars*, (E): 100 μm, (F): 50 μm.

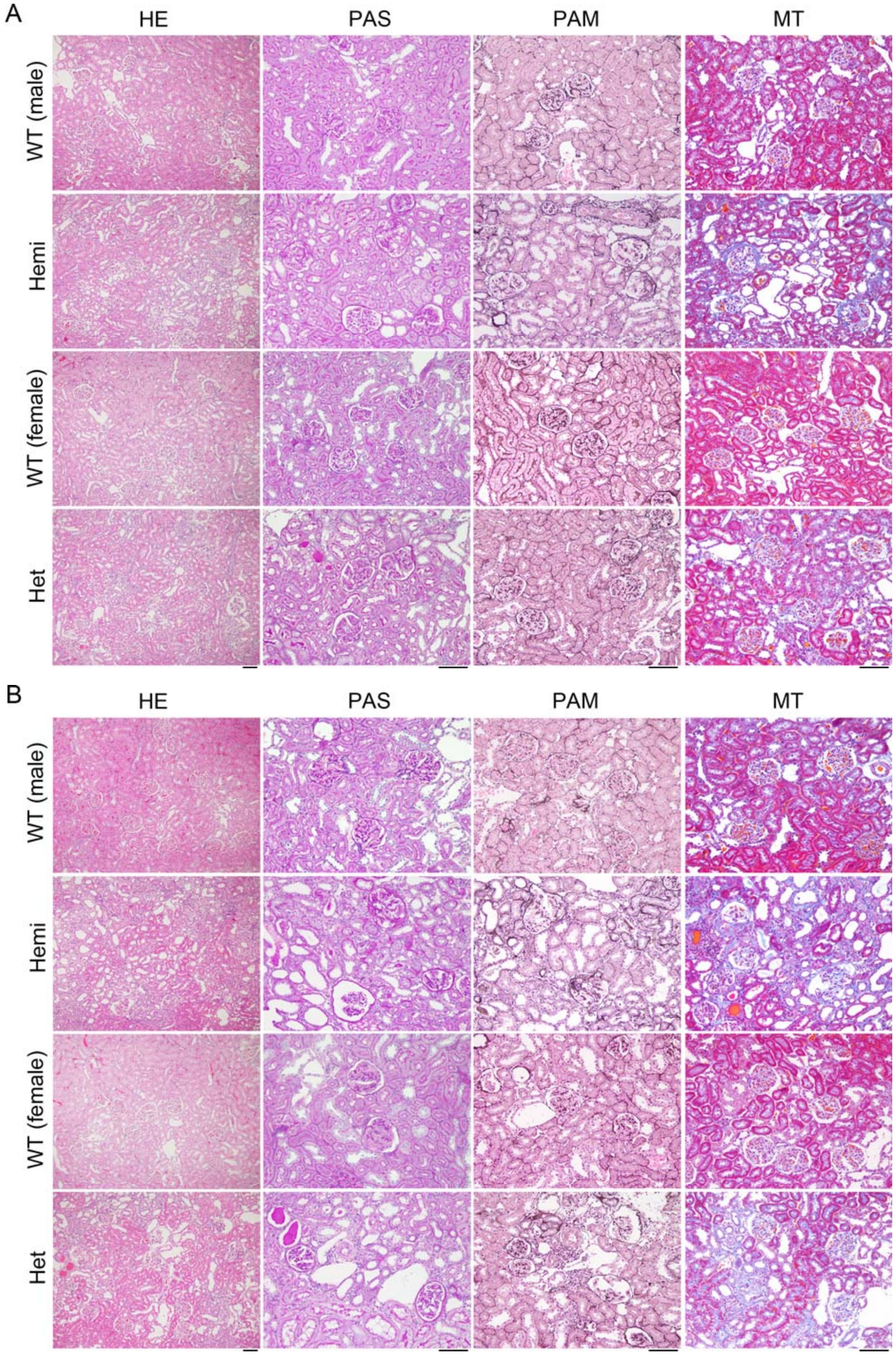


**Supplemental Figure 4. Histological analyses of the *Col4α5* deficient kidneys**
Representative microscopic images in wildtype (WT) and *Col4*α*5* mutant (Hemi; hemizygous males, Het; heterozygous females) rats at 12 weeks (A) and 16 weeks (B) of age. These tissue sections were prepared and stained with hematoxylin and eosin (HE), Periodic acid Schiff (PAS), periodic acid methenamine silver (PAM), and Masson trichrome (MT). *Scale bars*, 100 μm.

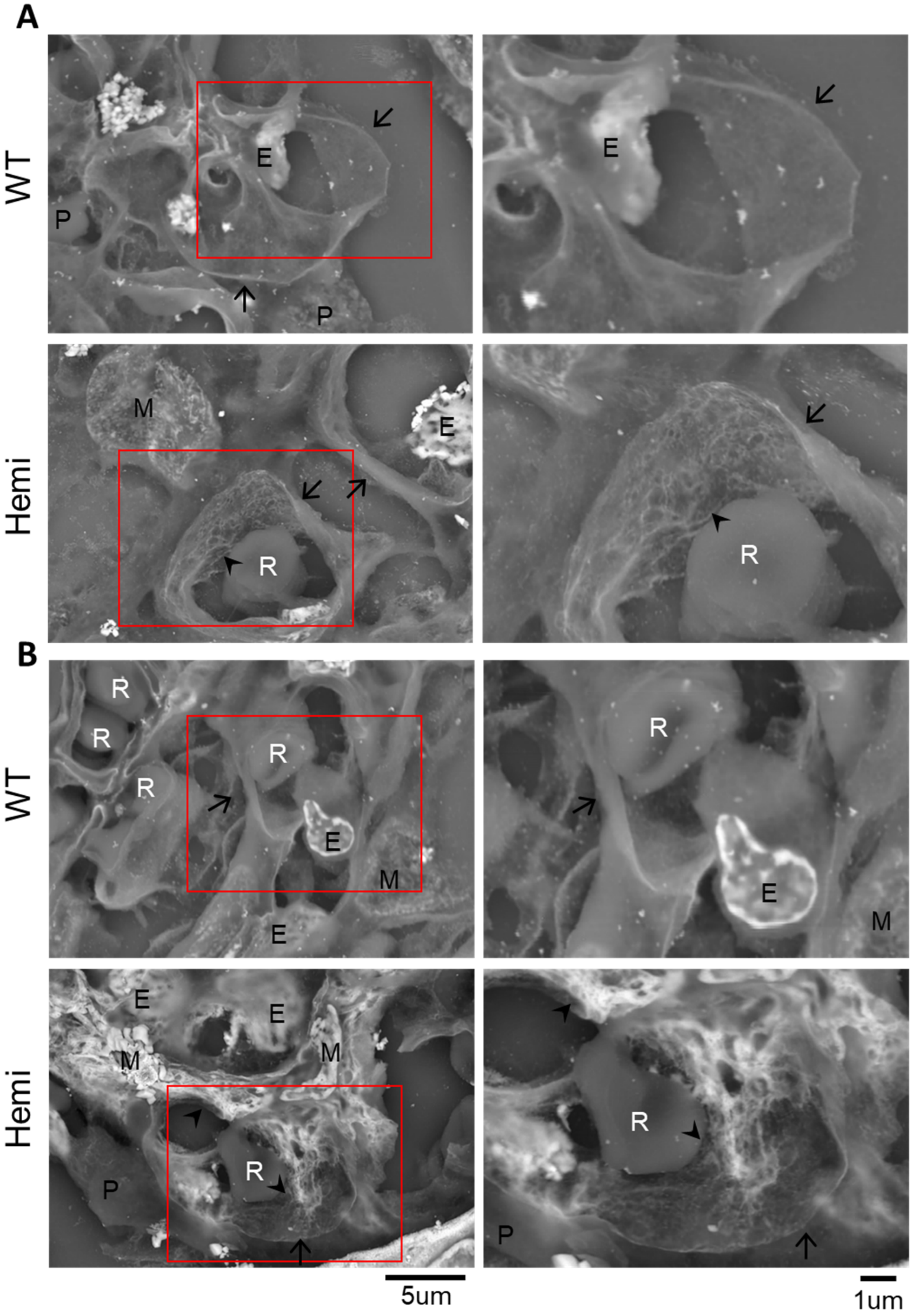


**Supplemental Figure 5. Electron photomicrographs of glomerular basement membranes in *Col4α5* mutant rats**

Representative Low-vacuum scanning electron microscopy (LVSEM) images in wildtype (WT) and *Col4α5* mutant (Hemi) male rats at 12 weeks (A) and 16 weeks (B) of age. Arrowheads indicate the coarse meshwork structure of the GBMs. Arrows also indicate cut side of the capillary walls. Red insets are revealed the higher magnification of left panels. E: Endothelial cells, M: Mesangial cells, P: Podocytes, R: Red blood cells. *Scale bars*, 5 µm (left), 1 µm (right).

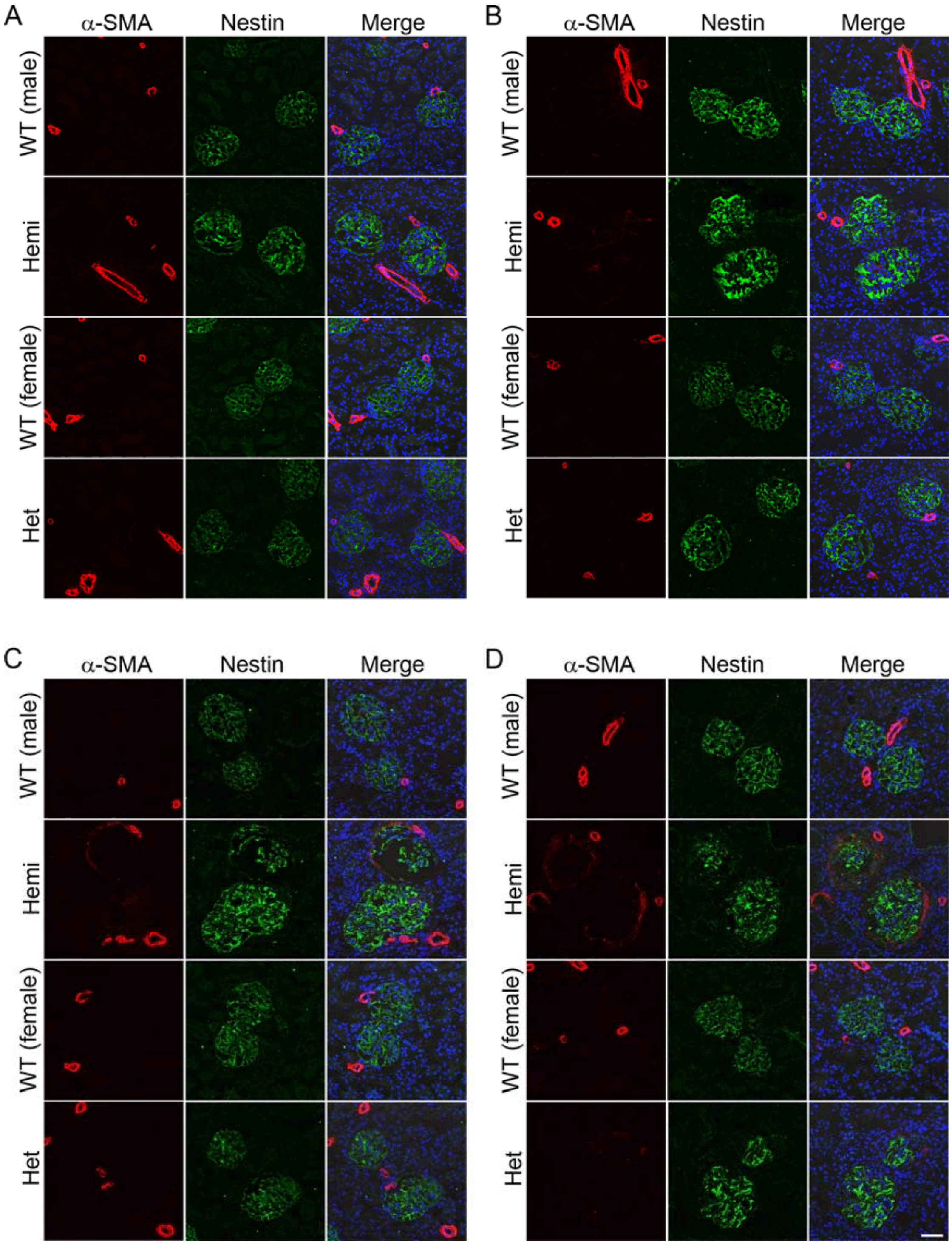


**Supplemental Figure 6. Renal fibrosis in *Col4α5* deficient rats**
(A-D) Immunostaining of kidney sections with α-SMA (red), nestin (green; glomeruli), and DAPI (blue; nuclei) in wildtype (WT) and *Col4α5* mutant (Hemi; hemizygous males, Het; heterozygous females) rats from 8 (A), 12 (B), 16 (C), to 20 (D) weeks of age. *Scale bars*, 50 μm.

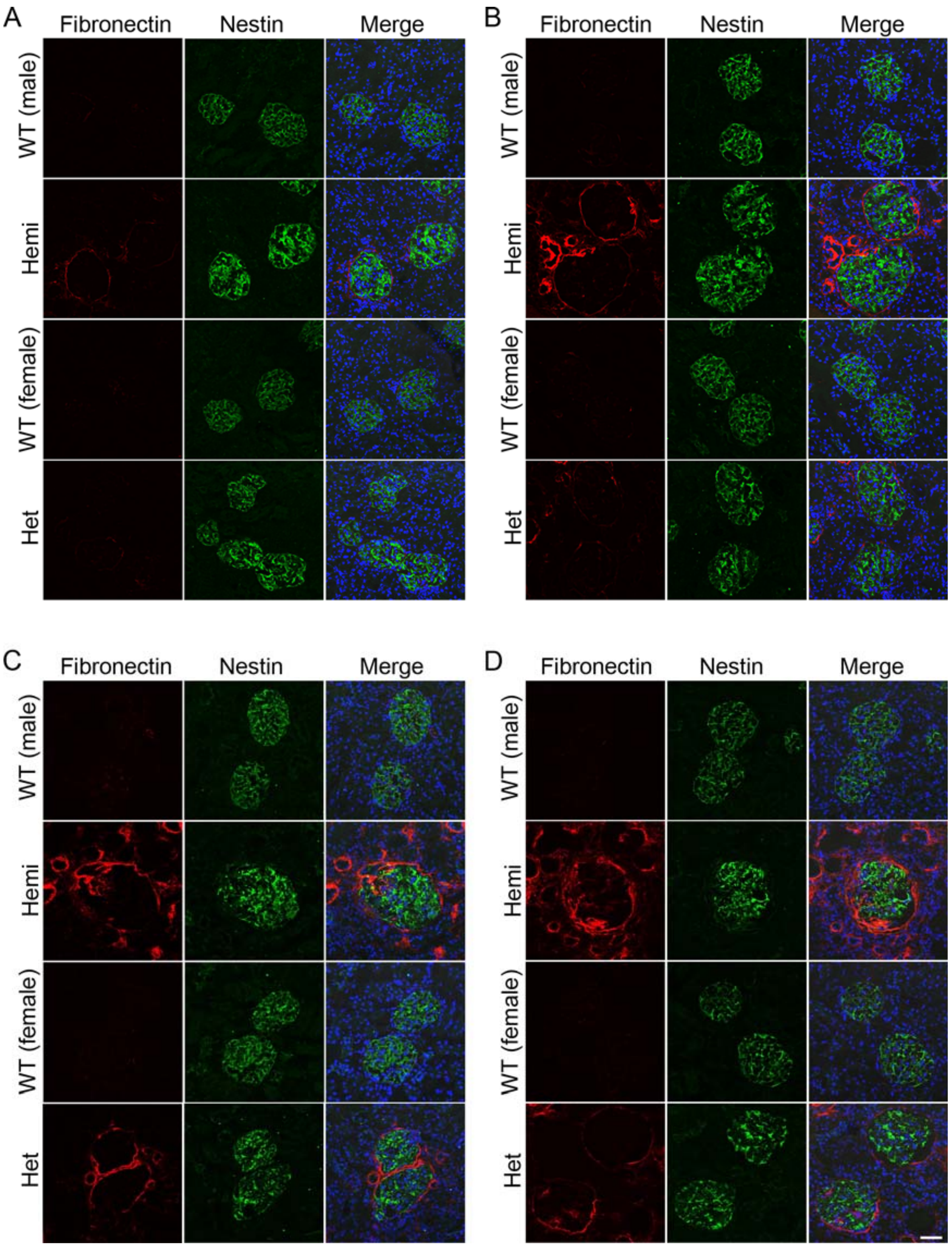


**Supplemental Figure 7. Renal fibrosis in *Col4α5* deficient rats**
(A-D) Immunostaining of kidney sections with fibronectin (red), nestin (green; glomeruli), and DAPI (blue; nuclei) in wildtype (WT) and *Col4α5* mutant (Hemi; hemizygous males, Het; heterozygous females) rats from 8 (A), 12 (B), 16 (C), to 20 (D) weeks of age. *Scale bars*, 50 μm.

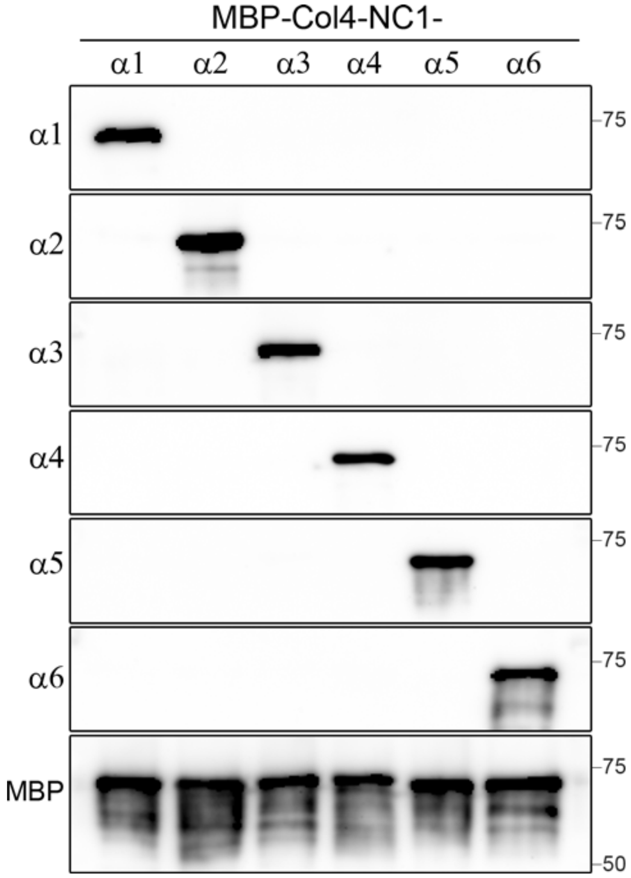


**Supplemental Figure 8. Characterization of an antibody specifically recognized type IV collagen**

Immunoreactivity was observed specifically with COL4A6 (α6), but not with other COL4 protein (α1-5). The type IV collagen α1-5 protein antibodies were also immunoreacted specifically with COL4 protein (α1-5) proteins, respectively.

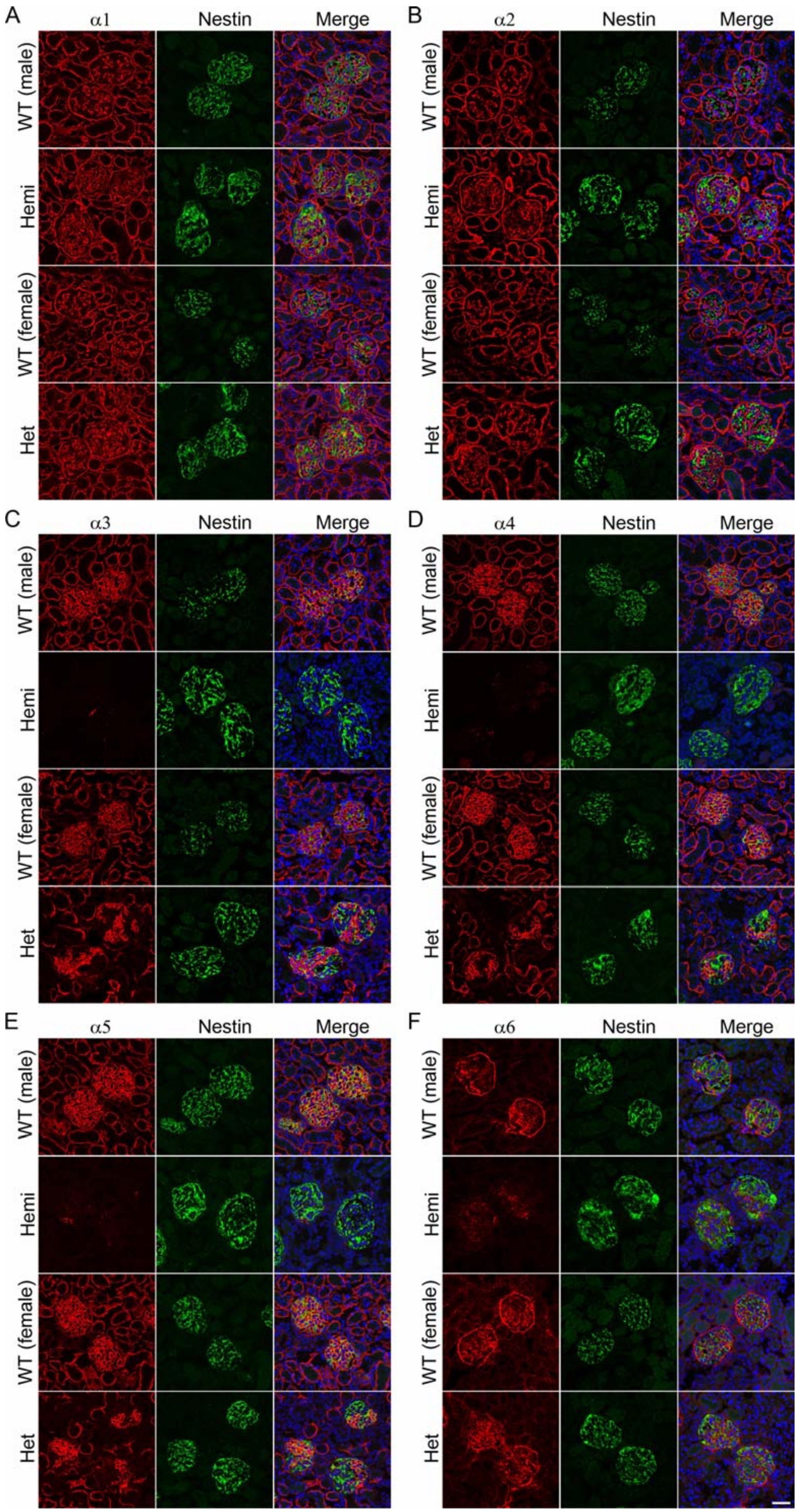


**Supplemental Figure 9. Type IV collagen distributions in the *Col4α5* deficient kidneys at 8 weeks of age**

(A, B) Immunofluorescence analyses of kidney sections with antibodies against α1-6 (IV) (red), nestin (green; glomeruli), and DAPI (blue; nuclei) in wildtype (WT) and *Col4α5* mutant (Hemi; hemizygous males, Het; heterozygous females) rats. *Scale bars*, 50 µm.

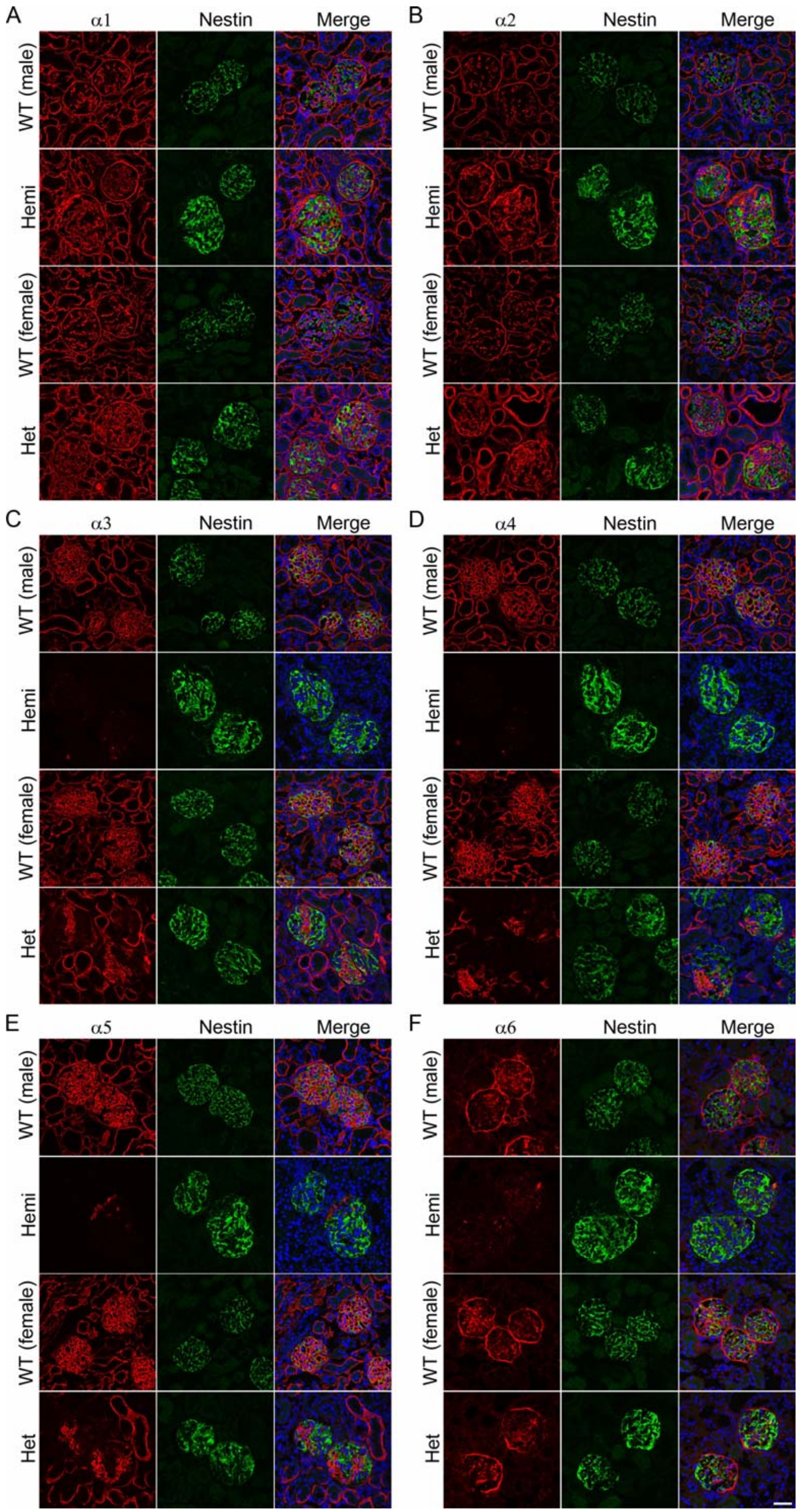


**Supplemental Figure 10. Type IV collagen distributions in the *Col4α5* deficient kidneys at 12 weeks of age**

(A, B) Immunofluorescence analyses of kidney sections with antibodies against α1-6 (IV) (red), nestin (green; glomeruli), and DAPI (blue; nuclei) in wildtype (WT) and *Col4α5* mutant (Hemi; hemizygous males, Het; heterozygous females) rats. *Scale bars*, 50 μm.

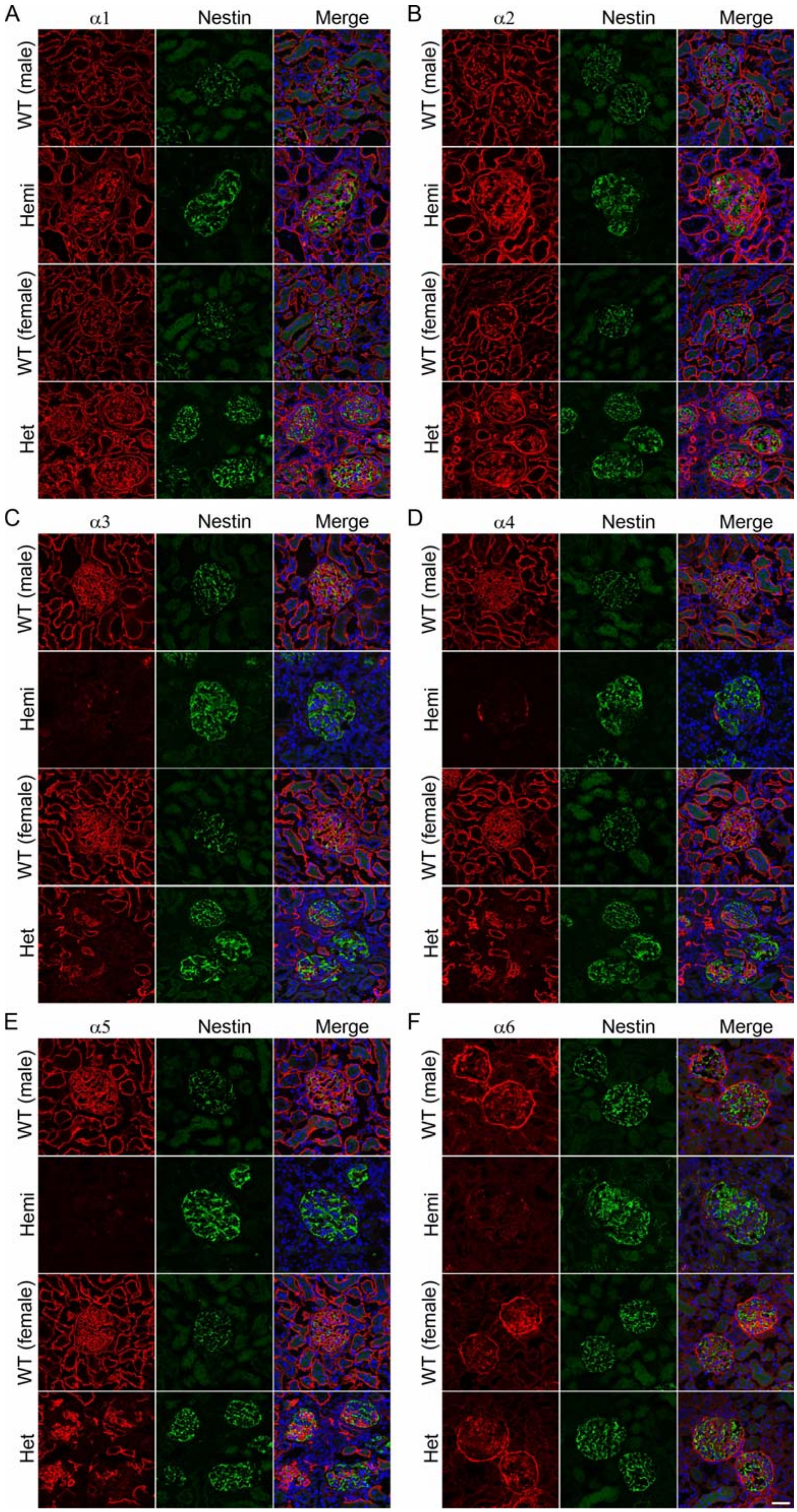


**Supplemental Figure 11. Type IV collagen distributions in the *Col4α5* deficient kidneys at 16 weeks of age**

(A, B) Immunofluorescence analyses of kidney sections with antibodies against α1-6 (IV) (red), nestin (green; glomeruli), and DAPI (blue; nuclei) in wildtype (WT) and *Col4α5* mutant (Hemi; hemizygous males, Het; heterozygous females) rats. *Scale bars*, 50 μm.

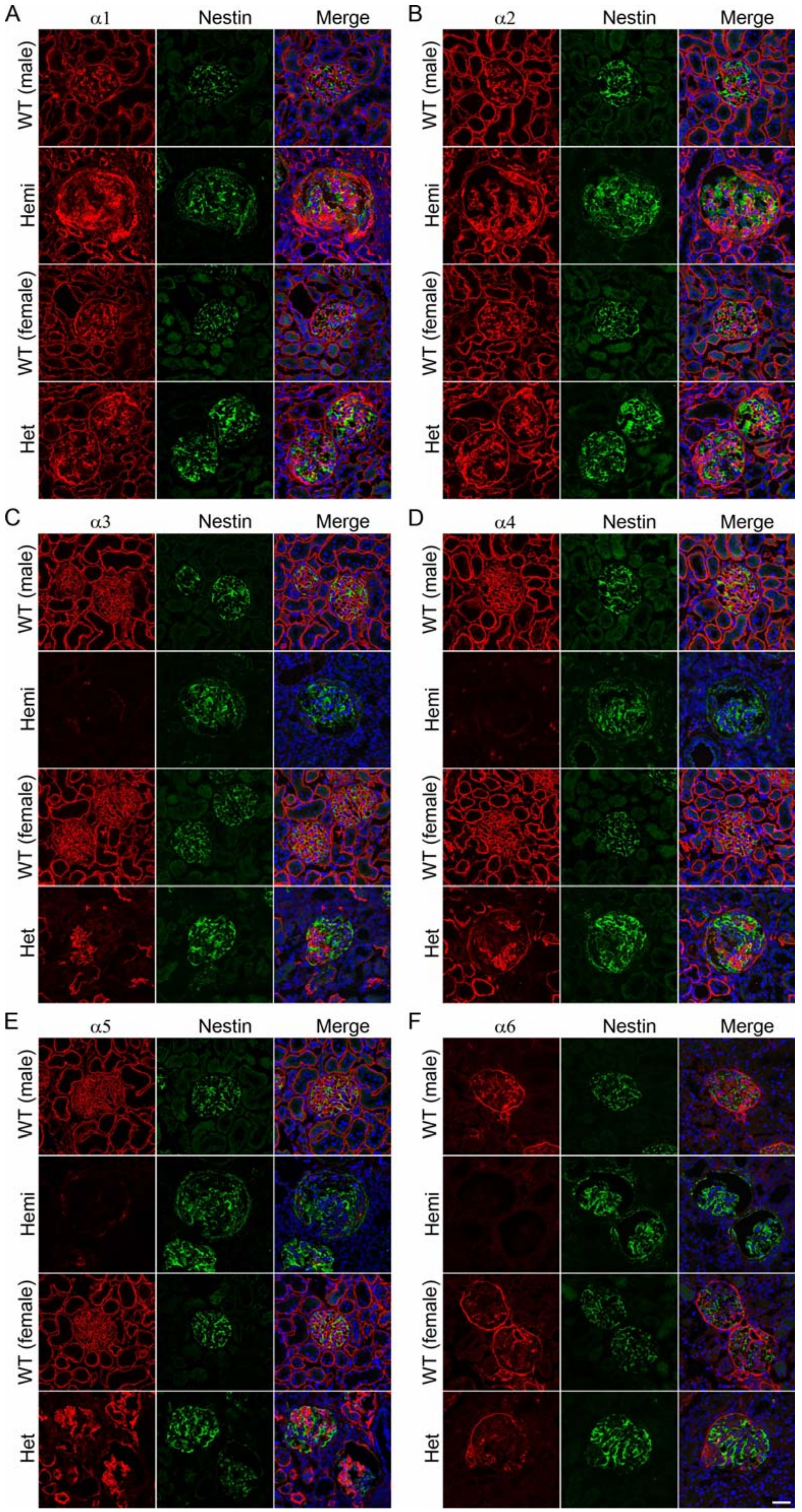


**Supplemental Figure 12. Type IV collagen distributions in the *Col4α5* deficient kidneys at 20 weeks of age**

(A, B) Immunofluorescence analyses of kidney sections with antibodies against α1-6 (IV) (red), nestin (green; glomeruli), and DAPI (blue; nuclei) in wildtype (WT) and *Col4α5* mutant (Hemi; hemizygous males, Het; heterozygous females) rats. *Scale bars*, 50 μm.

**Supplemental Table 1. Proteinuria in *Col4a5* deficient rats**

| | WT (male) | | Hemi | | WT (female) | | Het | |
|---|---|---|---|---|---|---|---|---|
| | proteinuria (mg/16h) | n | proteinuria(mg/16h) | n | proteinuria (mg/16h) | n | proteinuria(mg/16h) | n |
| 4W | 0.1 ± 0.0 | 10 | 0.6 ± 0.1 | 16 | 0.1 ± 0.0 | 12 | 0.5 ± 0.1 | 11 |
| 6W | 0.4 ± 0.1 | 10 | 6.2 ± 0.4 | 12 | 0.4 ± 0.1 | 8 | 2.0 ± 0.5 | 10 |
| 8W | 2.7 ± 0.2 | 32 | 16.4 ± 0.9 | 57 | 0.7 ± 0.0 | 58 | 3.5 ± 0.3 | 42 |
| 12W | 9.2 ± 0.7 | 35 | 77.0 ± 3.8 | 59 | 1.1 ± 0.0 | 59 | 14.2 ± 1.8 | 43 |
| 16W | 8.9 ± 0.7 | 37 | 87.5 ± 6.8 | 59 | 1.2 ± 0.1 | 60 | 30.5 ± 3.8 | 45 |
| 20W | 8.5 ± 0.5 | 35 | 100.1 ± 12.9 | 53 | 1.1 ± 0.0 | 58 | 42.9 ± 5.3 | 41 |
| 24W | | | | | 1.1 ± 0.0 | 58 | 57.8 ± 7.8 | 40 |
| 28W | | | | | 1.0 ± 0.0 | 57 | 85.3 ± 10.4 | 39 |
| 32W | | | | | 1.0 ± 0.0 | 57 | 87.3 ± 10.9 | 39 |
| 36W | | | | | 1.0 ± 0.0 | 51 | 124.1 ± 12.4 | 36 |
| 40W | | | | | 1.0 ± 0.1 | 49 | 133.8 ± 15.9 | 34 |
| 44W | | | | | 1.1 ± 0.1 | 47 | 157.2 ± 25.0 | 32 |
| 48W | | | | | 1.2 ± 0.1 | 40 | 163.2 ± 30.5 | 30 |
| 52W | | | | | 1.3 ± 0.1 | 40 | 175.7 ± 22.0 | 30 |

wildtype (WT), Hemizygous (Hemi), and Heterozygous (Het) mutant rats
